# Developing and Benchmarking One Health Genomic Surveillance Tools for Influenza A Virus in Wastewater

**DOI:** 10.1101/2025.09.19.676942

**Authors:** Minxi Jiang, Audrey L.W. Wang, James B. Thissen, Kara L. Nelson, Lenore Pipes, Rose S. Kantor

**Author notes:** Corresponding author: Rose S. Kantor, Lenore Pipes.

## Abstract

Influenza A viruses (IAV) remain a persistent One Health threat, and whole-genome sequencing from wastewater offers a promising surveillance tool. However, IAV is at low abundance in wastewater, making it difficult to sequence. We benchmarked four targeted enrichment methods suited for whole-genome sequencing including custom and off-the-shelf amplicon and probe-based methods. Our custom HA tiled-amplicon panel was sensitive, fast, and cost-effective, making it suitable for monitoring low-abundance seasonal variants of known subtypes. However, its reliance on conserved and intact primer-binding sites limited primer design to fewer subtypes. A previously published universal amplicon method targeted all IAV subtypes, but it performed poorly in wastewater due to its reliance on intact genome segments. Probe-capture methods were resilient to RNA degradation and mismatches, potentially enabling broader surveillance and detection of emerging strains. However, probes were costly, labor-intensive, and less sensitive than tiled-amplicon. When testing compatibility of sequencing methods with upstream virus concentration and extraction methods, ultrafiltration-based virus concentration outperformed large-volume direct extraction with all four sequencing methods. This set of benchmarking comparisons and custom panels provides needed information for the translation of IAV genomic sequencing into a routine component of wastewater surveillance.

## 1. Introduction

Influenza A viruses (IAV) are panzoonotic, and animal spillover events have led to at least 5 previous pandemics in the 20th and 21st centuries^1,2^. Of recent concern, highly pathogenic avian influenza H5N1 clade 2.3.4.4.b reached wild birds in the US in 2022, subsequently spreading to poultry, dairy cattle, and humans with multiple species crossover events since 2024^3^. In this context, whole genome sequencing of human and animal specimens has provided critical information for tracking the origins of human infections, detecting adaptation to human hosts and antiviral resistance, and examining the suitability of existing candidate vaccine viruses^4^. Meanwhile, the US CDC conducts routine monitoring of human seasonal influenza through whole genome sequencing of approximately 7,000 clinical specimens each year. These sequences are used to inform vaccine development for seasonal IAV strains (H3N2 and H1N1)^5^. However, experiences from surveillance during the COVID-19 pandemic highlighted biases in clinical surveillance systems due to differences in symptoms, testing availability, and test-seeking behavior for different populations^6^. Wastewater-based genomic surveillance has the potential to complement clinical data and to integrate human and animal testing with fewer biases, operationalizing the One Health paradigm.

Acquisition of high-quality influenza A whole genome sequences from wastewater is challenging due to viral genetic diversity across multiple subtypes, the segmented nature of the genome, low concentrations of target viral RNA, and a complex sequencing background. To overcome these challenges, three principal enrichment strategies have been applied, each with limited success: first, a whole genome amplification method using universal primers was paired with nanopore sequencing, making use of the highly conserved 5’ and 3’ terminal sequences present on all segments across subtypes^7^. Lee et al. reported sequencing data for just 12 of 48 samples tested, and no single sample yielded all eight segments^8^. Second, several multi-virus probe-capture panels have also been applied to detect the whole genome of IAV in wastewater, including Illumina’s RVOP^9–11^, Qiagen’s xHYB adventitious agent panel^12^, and Twist’s Comprehensive Viral Research panel^13^. However, in all reports, IAV genomes were highly incomplete unless data from many samples were combined. While whole-genome recovery is valuable for identifying reassortment events and mutations across different segments, each carrying different public health risks, prioritizing key segments may offer a faster and potentially more sensitive alternative for subtype tracking, vaccine development, and early warning. A recently developed tiled amplicon panel covering the HA, NA, and M segments of seasonal subtypes was applied to wastewater, with successful sequencing of H1N1 but limited detection of H3N2^14^. A separate study improved H3N2 coverage by using shorter amplicons, achieving high-quality HA segment recovery at sufficiently high viral concentrations^15^. Commercial tiled amplicon panel targeting H1N1, H1N2, and H3N2 (CleanPlex Respiratory Virus, Paragon Genomics) was also applied to wastewater, yielding high recovery for six of eight IAV segments^16^.

The quality of extracted nucleic acids from wastewater may also limit IAV genome recovery. RNA degradation during wastewater transit and sample processing could lead to fragmented genomes that cannot be enriched by certain downstream sequencing methods^17^. Reflecting this challenge, we previously found that wastewater processing methods that were more sensitive for PCR-based viral detection did not necessarily yield the best results for sequencing^18^, Additionally, we predicted that the ratio between target viral nucleic acid and non-target nucleic acids in a sample may affect the sensitivity of probe-capture sequencing. This ratio is largely determined by the method of wastewater concentration and by the number of targets in the probe panel^19^. Further work is needed to understand the interaction between sample processing methods and sequencing methods for specific viruses of interest such as IAV.

The goal of this study was to develop a One Health-focused, wastewater-based genomic surveillance workflow, which is capable of sensitively, efficiently, and cost-effectively detecting and characterizing both circulating and emerging influenza A viruses (IAVs). To our knowledge when we embarked on this research, no publicly available tiled amplicon methods yet existed for IAV subtype H5N1. A more recently published study has since developed a tiled-amplicon approach targeting all eight segments of H5N1, but only for cattle-associated strains^20^. Similarly, no IAV-specific probe set could target both zoonotic and seasonal human subtypes, although human-specific broad viral panels (Twist CVRP, Illumina VSP) and avian-specific IAV panels existed^21^. This motivated us to design two custom panels (1) a HA-segment tiled-amplicon panel prioritized for H1N1, H3N2, and H5N1 subtypes; (2) a probe-capture panel covering the whole genome (8 segments) of 11 representative panzoonotic IAV subtypes. We benchmarked the performance of these assays against two existing sequencing methods (**Table 1**): (3) a pan-viral whole-genome probe-capture panel (Probe-Twist), and (4) a whole-genome amplification method covering all IAV subtypes and segments (Universal-amplicon)^7^. We report on challenges in custom panel design, sequencing sensitivity, interactions with upstream virus concentration methods, and logistical factors such as cost and turnaround time.

**Table 1.** Sequence enrichment methods applied in this study.

| Method | Targeted segment /subtype | Sequencing | Input vol. (uL) | PCR cycles | Insert size (bp) | Avg. Depth per sample (M reads)* | Source |
| --- | --- | --- | --- | --- | --- | --- | --- |
| Tiled-amplicon | HA / H1N1, H3N2, H5N1 | NextSeq 300 PE | 5 | 35+8 | 400-565 | $8.4 \pm 1.5$ | This study |
| Probe-IAV | All / 11 subtypes** | NextSeq 300 PE | 15 | 12+17 | 300-500 | $20.3 \pm 2.0$ | This study |
| Probe-Twist | All / 119 | NextSeq 150 PE | 15 | 10+8 | 180-220 | $2.8 \pm 0.1$ | Twist |
| Universal-amplicon | All / all | minION | 5 | 35 | | $0.5 \pm 0.2$ | Zhou *et al.* (2009)<sup>7</sup> |
| Unenriched | All / all | NextSeq 300 PE | 15 | 12 | 300-500 | 14.4 | NA |

\* All values are reported for S1, except for the unenriched samples, for which the sequencing depth is reported for S3. The sequencing depths for other samples are included in **Table S7**.
\*\*see Supplementary Methods.

## 2. Results

### 2.1 Design of the Probe-IAV and HA segment tiled-amplicon panel

To compile a dataset for probe design, we first explored sequences and host diversity of IAV genomes. Searching GISAID for the 11 most prevalent panzoonotic IAV subtypes yielded 486,782 complete genomes collected within the previous year (**Supplementary Methods**). We noted that the number of H1N1 and H3N2 sequences released in a single year exceeded the total 5-year data for H5N1 and all archived sequences of other subtypes (**Table S1**). Further, the recent H1N1 and H3N2 sequences were predominantly from human hosts. To improve host diversity, we added avian and swine H1N1 and H3N2 sequences from the past 10 years (**Figure S1**). We also included sequences from the four spike-in strains used for benchmarking, resulting in a final dataset of 525,075 sequences. We used Syotti for probe design, testing multiple strategies to balance genome coverage, probe count, and hybridization efficiency across IAV genomes. Clustering input sequences at 100% identity initially produced up to 69,835 probes. A stepwise design approach, which targeted 90% of genomes with stricter mismatch limits and the remaining 10% with relaxed criteria (e.g., allowing gaps), helped reduce the probe count but still exceeded our target of fewer than 7,500 probes (**Table S2**). Ultimately, clustering IAV genomes at 90% identity resulted in a final set of 7,448 probes (Design 15, **Table S2**). When probes were aligned to the spike-in reference genomes, all spike-in viruses showed coverage breadth >75%, except for H3N1, where the MP and PB2 segments exhibited lower coverage breadth (**Figure S2**). We also observed regions with higher predicted probe density (>4x), particularly at the ends of each segment in the H1N1 genome.

Tiled primer design used a dataset consisting of HA sequences from the previous one-year for H3N2 and H1N1, and 5 years for H5N1. No geographic or host-based restrictions were applied during sequence selection. Unlike probe design, clustering was performed at 100% identity to account for the higher specificity of PCR. The final datasets contained sequences from 8558, 5876, and 6052 strains for H1N1, H3N2, and H5N1, respectively. We investigated design tools including PriMux^22,23^, PrimalScheme^24^, and Olivar^25^. The latter two tools have since been updated, but at the time of our design, PrimalScheme input was limited to 200 reference genomes, while Olivar required a risk profile rather than an alignment. No tool could accommodate segmented genomes as input. These restrictions prompted us to choose PriMux and to focus solely on the HA segment to limit complexity. Although the kmer-based-approach of PriMux allows amplification of a broader diversity of sequences, we chose to perform separate runs for each subtype to improve specificity. PriMux produced shorter-than-requested amplicons, and after tuning parameters, the resulting amplicons were typically around 400 nt. However, some amplicons were shorter or longer than expected and/or spanned other amplicons. Manual review was required for several steps, resulting in a lengthy design process: first, more than one primer set per amplicon was often produced, and manual review was required to choose the best one; second external tools were required to predict primer dimers and hairpins, and the melting temperatures of the primers varied widely; third, compatibility between primer sets from multiple subtypes required manual investigation; lastly, we found some cases where none of the PriMux primer options for a given tile were suitable based on our review, necessitating manual redesign (see **Supplementary Methods**).

### 2.2 Comparison of coverage breadth and depth among different methods for IAV sequencing

Four targeted IAV sequencing methods (**Table 1**) were benchmarked by sequencing three RNA mixtures in triplicate. The mixtures, “S1”, “S2”, and “S3”, were prepared by combining purified RNA from four IAV strains (subtypes H1N1, H3N2, H3N1, and H5N1) with IAV-negative wastewater RNA (**Figure 1a**). The resulting target-to-background RNA mass ratio increased by approximately 100-fold from S1 to S3, (4.4×10⁻⁸ to 2.4×10⁻⁶), although for H3N1 and H5N1, equal concentrations were targeted to spike into mixtures S2 and S3 due to limited viral titers. Unenriched metagenomic sequencing was also performed with mixture S3 to assess the fold-enrichment of the targeted sequencing methods.

**Figure 1.**
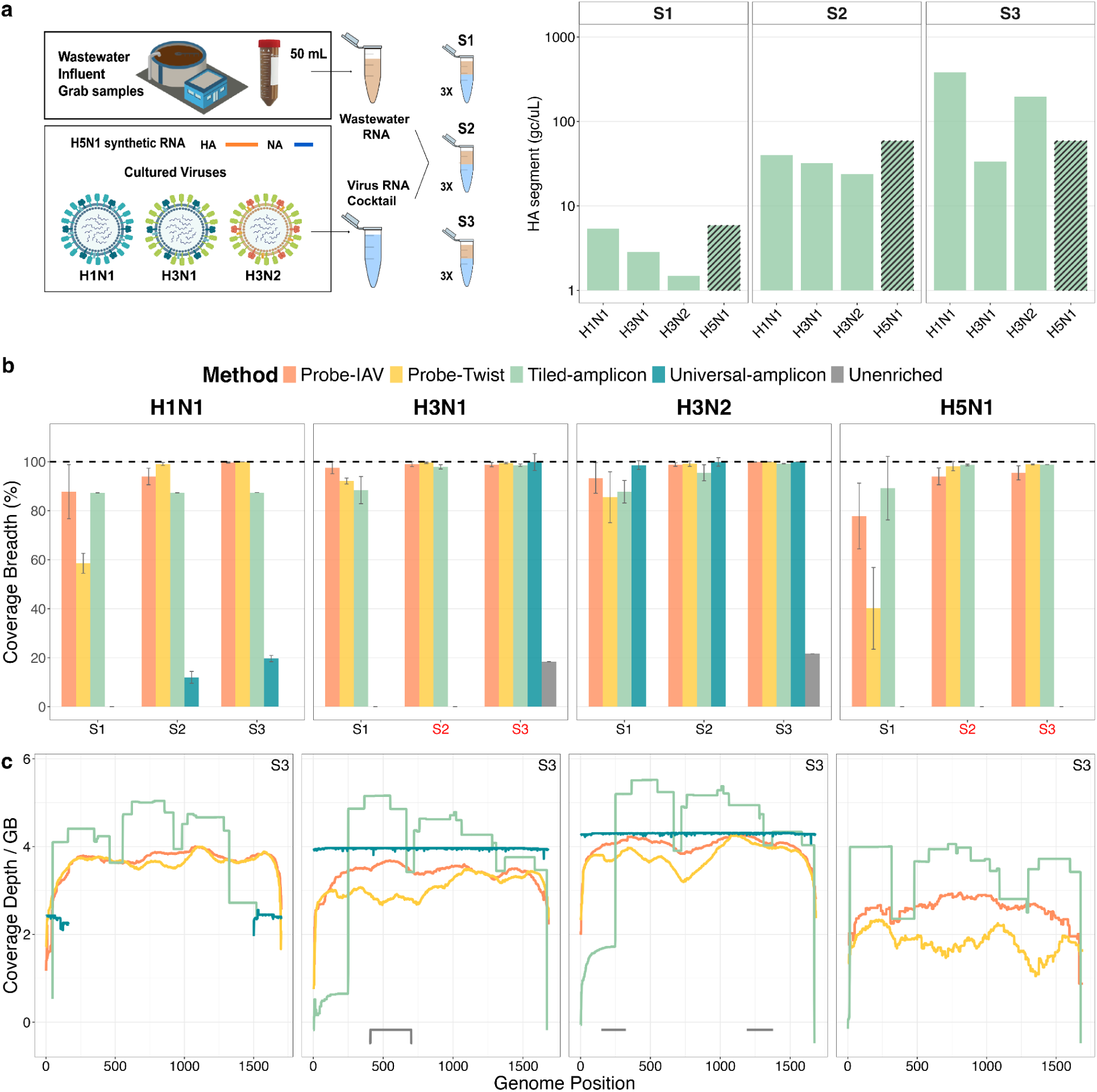
Performance of different sequencing methods for detecting the HA segment of IAV. **(a)** Schematic showing the preparation of the three RNA mixtures and the actual concentrations of each strain in each mixture measured by dPCR. Note that for H3N1 and H5N1, equal concentrations were added to mixtures S2 and S3 due to limited viral titers. Striped bars of H5N1 indicated the target concentrations calculated from dPCR-validated stock measurements rather than concentrations directly measured in mixture S1 to S3. **(b)** HA segment coverage breadth produced by each method, for each strain within samples S1, S2, and S3 (n=3 replicates per sample). H3N1 and H5N1 were spiked at the same concentration in S2 and S3 (red labels). **(c)** HA segment coverage depth (log₁₀ depth at each position per total Gbp sequenced) in the highest spike-in sample (S3), shown for each method and strain.

Across all sequencing methods, the coverage breadth of HA segments increased with increasing viral RNA concentrations. At the lowest concentration (mixture S1), the Universal-amplicon method recovered the HA segment only for H3N2, while Probe-Twist panel recovered 86 ± 10% for H3N1 and H3N2, but only 59 ± 4% for H1N1 and 40 ± 17% for H5N1 (**Table S8**). The two customized approaches, Tiled-amplicon and Probe-IAV, achieved over 78 ± 13% coverage for all four strains (**Table S8**). At the highest IAV concentrations (mixture S3), Tiled-amplicon, Probe-IAV, and Probe-Twist each recovered near-complete HA segments for H3N1, H3N2, and H5N1 (coverage breadth >98.5%, **Figure 1b, Table S8**), while tiled amplicon coverage of H1N1 remained limited to 87% due to one failed tile that requires further PCR optimization. Universal-amplicon did not produce reads for HA from H5N1 at any concentration, as the synthetic RNA lacked the primer-binding regions. Notably, all enrichment methods outperformed unenriched metagenomic sequencing, which recovered less than 22% of the HA segment for H3N1 and H3N2 even at the highest input concentration (**Table S8**). We note that coverage breadth, or the fraction of bases in the genome that are covered by sequencing reads, is somewhat dependent on the total number of reads sequenced, as well as the sequencing platform, insert size, and read length. As these factors differ across methods, the comparison of coverage breadth limited to each method exactly as it was applied in this study. To assess potential regional biases in HA segment recovery, coverage depth profiles for each method were examined in mixture S3 (**Figure 1c**). The Tiled-amplicon method generally showed higher normalized coverage depth for most regions of the HA segment, and exhibited a repeating pattern of peaks and valleys due to tile overlap, along with reduced coverage for one tile of H3N1 and H3N2. Probe-IAV provided the most uniform HA segment coverage across all strains. In contrast, Probe-Twist exhibited coverage dropouts in H3N2 (500-1000bp) and H5N1 (1200-1500bp). The Universal-amplicon method recovered only the ends of the H1N1 HA segment, while unenriched samples recovered only short and randomly distributed fragments.

For the three whole-genome methods, we expanded our coverage analyses to include all eight segments (**Figure S3**). Here, the two probe-capture methods both performed well for mixture S3 (**Figure S3a** and **Table S8**). For mixture S1, both showed similar coverage breadth across different strains, but Probe-IAV provided higher and more even coverage depth than Probe-Twist (**Figure S3b**). Notably, when compared to the predicted coverage results, Probe-IAV showed higher empirical coverage breadth for segments with low predicted probe density (e.g., PB2 and NS of H3N1 showed over 95% coverage despite predicted recovery of less than 75%; **Figure S2a**). Additionally, regions with higher designed probe density did not consistently correspond to greater coverage depth; instead, coverage was generally uniform across the genome (**Figure S2b vs. Figure 1c**). Unlike the two customized methods, the universal-amplicon method continued to perform poorly for mixture S1, achieving good coverage across all strains only for the two shortest segments, MP and NS (> 99.4%, **Table S8**), while coverage for other segments varied widely among strains (**Figure S3a**).

### 2.3 Comparison of sensitivity and quantitativeness of different sequencing methods

To compare the sensitivity and potential enrichment bias towards different strains, we analyzed the relationship between library input quantity (as determined by dPCR assays targeting the HA segment of each strain) and the resulting coverage depth (RPKM) across all sequencing methods and strains (**Figure 2a**). H5N1 was excluded from this correlation because dPCR failed for H5 in spike-in samples, even though three sequencing methods yielded reads for H5N1 (**Figure 1a**).

**Figure 2:**
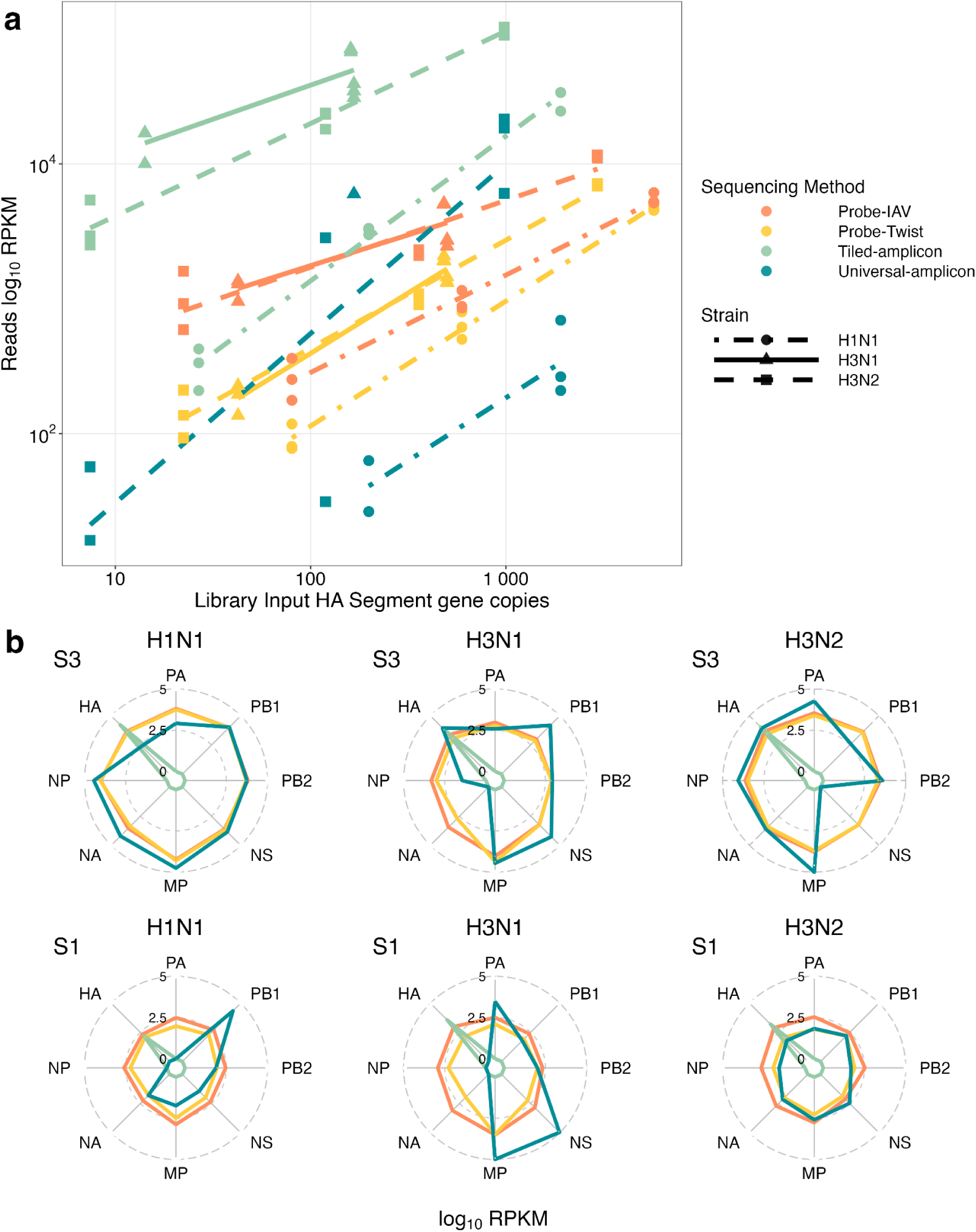
(a) Correlation between sequencing library input (HA segment copy number measured by dPCR) and the normalized coverage depth (RPKM) for each sequencing method. Note that library input gene copies accounts for differences in library input volumes. (b) log10(RPKM) values of all 8 segments within each virus strain, colored by sequencing method for mixture S3 (top) and mixture S1 (bottom). Note that in many cases, Probe-IAV and Probe-Twist are completely overlapping.

The correlations were strong for Probe-Twist across all strains (R² = 0.96-0.99, **Table S9**), while Universal-amplicon displayed the weakest correlations (for H1N1 and H3N2; R² < 0.82, **Table S9**). Tiled-amplicon and Probe-IAV performed well for H1N1 (R² ≥ 0.97). Notably, the slopes of the regression lines for different IAV subtypes were generally similar within sequencing methods, with highly parallel or overlapping trends (e.g., H3N1 and H3N2 in the two probe-capture methods, **Figure 2a**). However, regression lines for H3 and H1 were vertically offset within a given sequencing method, showing that normalized coverage depth is predicted to differ by subtype, even if library input quantities are held constant. Among all sequencing methods, the Tiled-amplicon was the most sensitive, yielding the highest coverage depths for all strains, particularly at the lowest input levels (< 10 gc). Universal-amplicon showed lower overall sensitivity but had a steeper response curve for H3N2 (slope = 1.25, **Table S9**), with RPKM values eventually surpassing that of Probe-Twist and Probe-IAV at higher input concentrations. The two probe-capture methods showed similar performance across the tested concentration range, but more gradual slopes for Probe-IAV suggest this method may be more sensitive at low inputs (**Figure 2a**). When the coverage depth analysis was extended to compare across segments within a strain, both probe-capture methods (Probe-IAV and Probe-Twist) yielded relatively uniform RPKM values across segments (**Figure 2b**). In contrast, the Universal-amplicon method exhibited extreme biases towards random segments (**Figure 2b**). Notably, both probe-capture methods displayed a modest bias toward the MP segment.

### 2.4 Impact of extraviral RNA decay and concentration/extraction methods on IAV sequencing

To evaluate the impact of concentration/extraction methods and extraviral RNA decay on sequencing outcomes, we spiked equal volumes (10 uL) of cultured H1N1, H3N2, and H3N1 into 40 mL raw wastewater aliquots, which equivalent to 37, 278, and 1.17 gc/mL, respectively, based on dPCR measurements of extracted viral stock RNA (**Table S10**). Immediately after spike-in (time 0), RNA was extracted from one set of aliquots using two different concentration/extraction methods: Innovaprep + Powerviral (IP) and the large-volume Promega Wizard Enviro TNA kit (PMG). The remaining aliquots were incubated for 3 hours at room temperature to reach equilibrium partitioning of viruses to solids and to allow potential decay of extraviral RNA^26^, followed by RNA extraction in triplicate using both methods (**Figure 3a**). IAV was quantified in extracted RNA using dPCR (**Table S11**) and sequenced using the four methods.

**Figure 3.**
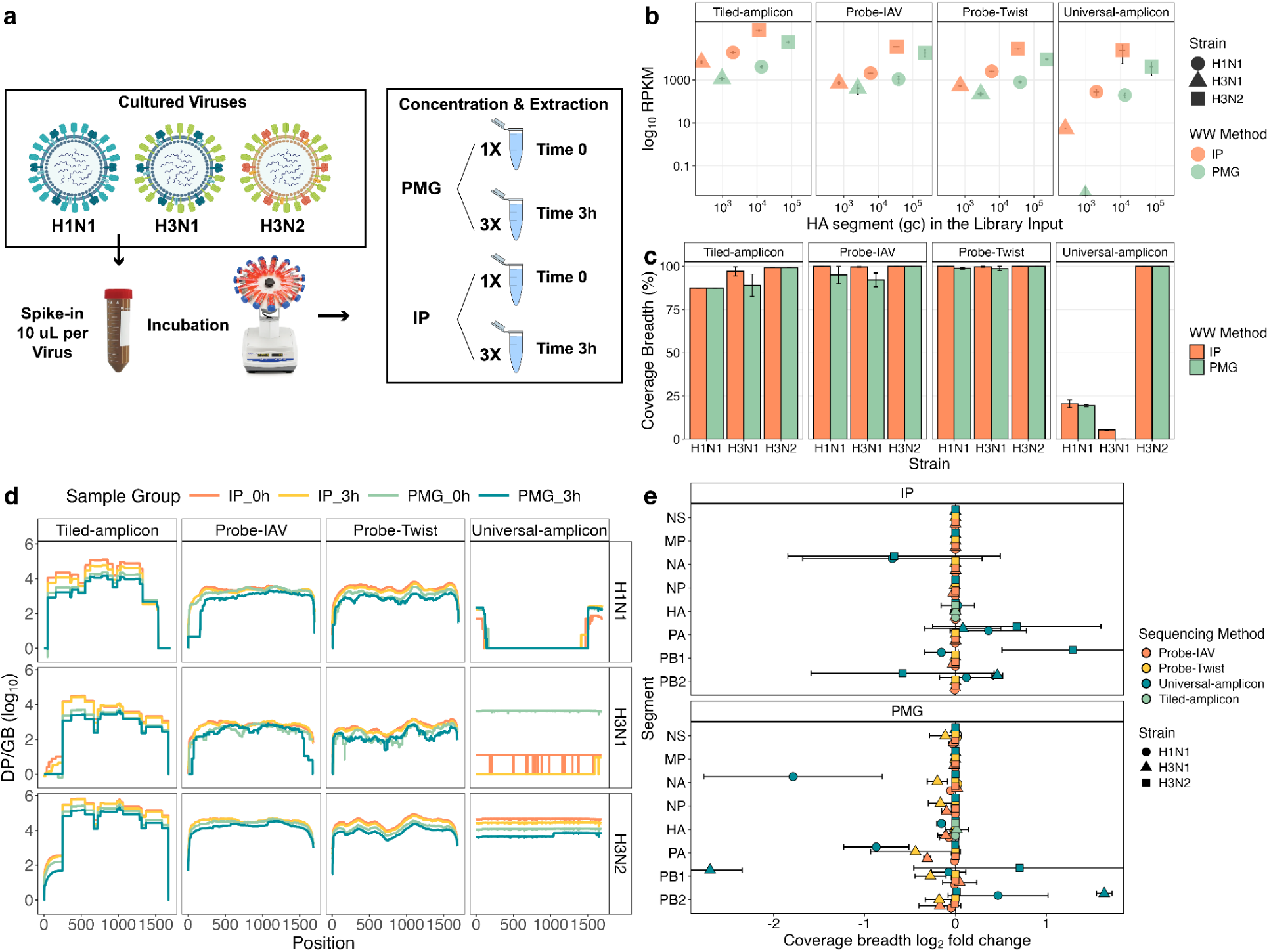
Comparison of concentration/extraction methods and the impact of incubation on genome-wide segment recovery across sequencing methods and IAV strains. (a) Experimental design illustrating virus spike-in, incubation, and concentration/extraction workflows. (b) Relationship between dPCR-measured IAV concentration in each sequencing library input and sequencing coverage depth (RPKM) for 3-hour incubated samples. Note that library input gene copies (gc) account for differences in library input volumes. (c) Relationship between wastewater concentration/extraction method, sequencing method, and HA segment coverage breadth for 3-hour incubated samples. (d) Coverage depth (log₁₀ depth per Gbp sequenced) across the HA segment before (0h) and after (3h) incubation, shown by strain and sequencing method. (e) log₂ fold-change of coverage breadth (%) before and after incubation of all eight genomic segments across four spike-in strains.

After three hours of incubation, the PMG method recovered significantly higher HA quantities (by dPCR) than IP for all IAV strains (**Figure 3b**, p < 0.05, **Table S12**). However, IP samples yielded higher normalized coverage depth, especially when sequenced with Tiled-amplicon and Probe-Twist (p < 0.05, **Table S12**). HA segment coverage breadth was similar between IP and PMG-treated samples, with >80% coverage across all strains when sequenced using Tiled-amplicon, Probe-IAV, and Probe-Twist. In contrast, coverage was generally poor with the Universal-amplicon method (**Figure 3c**). The potential impact of extraviral RNA decay on genome integrity was inspected by comparing the coverage across the HA genome. All methods except universal-amplicon consistently recovered the full HA segment across all strains, and no region-specific decay was observed. However, sharp drops in coverage depth were observed at the ends of the HA segment with the Probe-IAV method (first 100 bp for H1N1 and the last 100 bp for H3N1, **Figure 3d**). Overall, coverage depth generally followed the trend IP_0h > IP_3h > PMG_0h > PMG_3h across all sequencing methods, while Universal-amplicon showed greater variability (**Figure 3d**).

To assess the effects of decay across the whole genome, we compared the log₂ fold change (log₂FC) in coverage breadth between zero and three hours of incubation using both IP and PMG concentration/extraction methods (**Figure 3e**). IP samples showed little change in coverage breadth across virus strains and segments, with only minor decreases (log₂FC > 0.1) observed in Universal-amplicon. PMG samples exhibited greater coverage loss across more segments for all three whole-genome methods (**Table S13**). Universal-amplicon showed both increases and decreases in segment coverage with large standard deviations, which suggests stochastic amplification in PMG samples.

### 2.5 Economic analysis for different sequencing methods

To support practical decision-making and optimize wastewater processing workflows, we conducted a cost and labor analysis of the four sequencing methods (**Table S14**). The amplicon-based approaches had lower reagent costs ($173 per sample) and labor time (0.38 h per sample for H5 tiled-amplicon) compared to the probe-capture methods. The two probe-capture approaches had similar instrument and labor requirements, but the customized Probe-IAV was slightly less expensive than Probe-Twist in reagent costs ($533 vs. $572 per sample). Processing 24 probe-capture samples required ∼40 hours, nearly half of which was hands-off time for PCR cycling and hybridization (∼16-17 hours). In contrast, 57 tiled-amplicon samples were processed in ∼20 hours, with only ∼3 hours attributed to PCR cycling. The sequencing cost also varied depending on the required sequencing depth per sample. Although our study used the lowest sequencing depth for Probe-Twist due to cross-lab operational differences, in practice, the most sensitive method (tiled-amplicon) would likely incur the lowest sequencing depth cost.

## 3. Discussion

### 3.1 Performance of different targeted sequencing methods

The four benchmarked sequencing methods differed in their targets (HA segment vs. whole genome), reaction chemistry (PCR vs. hybridization), and library preparation protocols. Our analysis of these methods encompassed the complete end-to-end sequencing protocols tested here. Differences between the protocols resulted in varying sequencing performance, which can be leveraged to suit different surveillance objectives.

One objective is the ability to detect low-abundance emerging variants within the seasonal subtypes of IAV. The sensitivity (highest RPKM) and coverage breadth of the HA segment (>87% for all subtypes) achieved with tiled-amplicon make it well-suited for this application. Although our current panel targets only the HA segments of H1N1, H3N2, and H5N1 subtypes, the design principles and performance demonstrated here suggest that this approach can be extended to other segments, enabling broader, subtype-specific coverage as shown in recent studies^14,27^. An important concern is that viral evolution could lead to amplicon drop-out or decreased recovery, necessitating primer redesign^28^ (see below).

Another objective is whole-genome variant analysis of prevalent human and panzoonotic subtypes of IAV in wastewater. Such analysis is important to understand host range adaptation, antiviral resistance, and virulence^29,30^, and would be strengthened by the recovery of whole genomes for identification of unique mutations indicative of particular strains. In this regard, the Universal-amplicon method performed poorly, likely due to its dependence on high-quality, high-concentration genomic RNA inputs. (**Figure 1b, 1c, and 2b**). The two probe panels performed well, consistently recovering near-complete genomes. We observed differences at low input quantities (**Figure 1b, 1c, and 2b**), perhaps attributable to the probe design and sequencing settings: First, our custom Probe-IAV panel incorporated sequences updated through 2024, including H5N1 2.3.4.4b and all spike-in strains, whereas Probe-Twist used references only up to 2018, with the most recent H5N1 from 2017 (Casey Riegler, Twist Bioscience, personal communication, 7/11/25). Further, the broad viral target range of Probe-Twist may have reduced read coverage for low-abundance targets like IAV^19,31^. Second, due to differences in laboratory protocols, Probe-Twist had lower sequencing depth and shorter paired-read length than Probe-IAV (**Table 1**), which likely reduced its HA segment coverage breadth, especially in the lowest-concentration sample (S1). Despite these differences, the Probe-Twist panel still performed well, although with lower sensitivity for the H5N1 2.3.4.4b strain than the custom Probe-IAV panel. These results highlight the general robustness of the probe-capture approach to detect strains not included in the original design and suggest that regular design updates will increase sensitivity for emerging strains.

In addition to qualitative surveillance information, we were interested in the ability of sequencing to provide semiquantitative information about strain prevalence. This could be achieved if the sequencing method preserved relative abundance information. Our RNA mixtures contained a constant amount of wastewater RNA background with serially diluted viral RNA, but none of the sequencing methods reliably reflected the concentration ratios between subtypes as verified by dPCR (**Figure 2a**). Specifically, H1N1 was consistently undercounted relative to H3N1 and H3N2, while within subtypes, H3N1 and H3N2 showed less bias relative to each other (**Figure 2a**; highly overlapping lines). Such discrepancies are likely to become more complex in real wastewater samples, where viral composition varies longitudinally and geographically (Section 3.3). We recommend optimizing and balancing probes and primers based on their amplification and enrichment efficiencies across different subtypes, and integrating these with downstream deconvolution tools to improve quantitative accuracy in the future. In contrast to uneven subtype enrichment, the two probe-capture methods generally enriched all eight IAV segments evenly (**Figure 2b**), which might assist in identifying potential reassortments across different subtypes.

### 3.2 Design of Probe Capture vs. Tiled-amplicon Sequencing

We designed custom probe and tiled amplicon panels, allowing us to compare the ease of design and accuracy of *in silico* performance predictions for both. Successful design of broad panels was easier for probes than primers due to the availability of comprehensive design algorithms and to different constraints inherent in the enrichment mechanisms. In probe capture, random mismatches have been shown to be more detrimental than continuous mismatches^32^, leading us to choose a strategy that allowed gaps in some regions while keeping strict mismatch limits in others. *In silico* performance predictions by Syotti underestimated actual coverage breadth, likely because sequence mapping oversimplified these important details of hybridization chemistry.

Meanwhile, PCR-based enrichment is affected by mismatches only if they are in the primer-binding regions. Our simulate_pcr results predicted 2-3 primer mismatches which, although theoretically allowable under permissive conditions, caused certain tiles to fail *in vitro*. Additional design needs for PCR included consistent ranges of amplicon sizes and melting temperatures, and prevention of primer dimers and unwanted amplicons. These challenges would be expected to intensify when the number of primers increases, e.g., for panels targeting more genome segments, using shorter amplicons, or targeting increased diversity of IAV strains and subtypes. Recent primer design tools, such as Olivar model basic PCR reaction parameters^25^. Expanding models in tools like Olivar and Syotti to incorporate reaction dynamics and integrating iterative design-test-refine workflows could improve future performance. Overall, we found that increasing the diversity of input sequences increased both probe and primer counts in design outputs. While the cost of increased probes was higher compared to primers, the mechanistic constraints for increased primers were far greater than for probes.

Both probe-capture and tiled-amplicon panels may require updates over time because Influenza A viruses continually evolve through genetic drift and reassortment, which can introduce mutation-dependent detection bias as divergence accumulates^33^. In contrast, metagenomic sequencing is largely unaffected. The required update frequency of each method depends on the balance between the viral evolutionary rate and the method’s tolerance to mismatches. Probes generally tolerate fewer than five randomly distributed mismatches across the probe-binding region and can be easily updated with newly released sequences or genomes reconstructed from wastewater. However, the lead time and cost required would be expected to be higher than that of primers. Primers are more sensitive to mutations at binding sites, therefore, the frequency of primer updates for amplicon-based panels depends on whether the primer binding sites remain “low entropy” over time^25^. Designing primers that target broader and more conserved regions, where feasible within PCR constraints, could reduce the need for frequent redesign. For example, the Universal-amplicon panel is highly robust to viral evolution due to the conserved 5’ and 3’ termini in IAV genomes. As a practical recommendation, we suggest routinely mapping probe or primer sets to assembled sequences from wastewater, clinical and veterinary resources. This approach can help identify abnormal patterns, such as an unexpected decrease in detected reads, which may indicate the emergence of mutations and the need for probe or primer updates^34^. Lastly, although probe design updates are likely to be more straightforward, the lead time and cost required would be expected to be higher than that of primers.

### 3.3 Wastewater-Specific Challenges and Implications of One Health IAV surveillance

Real-world application of IAV targeted sequencing in wastewater must account for factors that affect RNA yield, integrity, and purity, including concentration and extraction methods, RNA decay in wastewater and freeze-thaw cycles. Based on our findings here and previously^18,35^, large-volume concentration and extraction (PMG) maximizes total IAV RNA yield, whereas ultrafiltration (IP) produces a higher proportion of IAV RNA relative to background, which is particularly beneficial for probe-capture methods where off-target reads are common (**Figure 3b**). IP also outperformed PMG for amplicon sequencing despite higher dPCR signals with PMG (**Figure 3b**). One plausible explanation is greater RNA fragmentation in PMG samples: longer amplicons require longer intact RNA segments containing both primer binding sites, making tiled-amplicon and especially universal-amplicon more vulnerable to fragmentation, whereas short dPCR targets remain amplifiable. However, spiking virus into wastewater prior to extraction does not allow viral RNA integrity to be distinguished from background wastewater RNA after extraction. This possible explanation will need to be explored using alternative approaches in future studies. We also caution that these differences between PMG and IP may be matrix-dependent and may not be consistently observed across broader wastewater conditions. Beyond extraction, viral RNA quantity and quality may also be impacted by RNA decay in wastewater or during freeze-thaw cycles after extraction. This could lead to either overall decreased virus signal or more fragmented RNA. We observed that wastewater incubation led to modest, genome-wide RNA degradation, affecting amplicon methods more severely due to their reliance on intact primer sites. Probe-capture showed a similar overall decline but with end losses and sporadic dips (**Figure 3d**). More experiments are needed to confirm whether wastewater processing causes region-specific degradation.

Ultimately, the choice of sequencing approach for IAV characterization in wastewater should be guided by the surveillance goal, available budget, and laboratory capacity. Practically, the tiled-amplicon assay is faster and more cost-effective to implement, making it well-suited for high-throughput subtyping of specific HA subtypes. In contrast, probe-capture is more labor-intensive and expensive (**Table S14**), but offers whole-genome coverage, broader subtype detection, and greater sensitivity in highly degraded samples.

The findings of this study have limitations when applied to real wastewater conditions. The spike-in samples used ranged from ∼50 to <5 genome copies/µL (S3-S1), which may not fully represent the lower and more variable concentrations typically found in wastewater^36^. Further, while our 3-hour spike-in incubation time likely accounted for adsorption kinetics and decay of extraviral RNA^26,37^, it may not fully capture viral decay that can occur during sewage transit and 24-hour composite sampling. Moreover, the use of a single grab sample of IAV-negative wastewater as the background does not account for the complexity and variability of inhibitory chemistry and microbiology across wastewater sources and over time. These factors may influence the observed quantitative relationship between dPCR and RPKM, especially for probe-capture methods. Further investigation is required to assess the performance of these sequencing methods under a range of real-world wastewater conditions.

## 4. Methods

### 4.1 Culturing and Acquisition of Spike-in Influenza A Viruses

Three IAV serotypes were obtained from BEI: H1N1 (A/California/04/2009), H3N2 (A/Netherlands/823/1992), and a reassorted H3N1 strain, containing the HA segment from H3N2 (A/Texas/1/77) and the remaining seven segments from H1N1 (A/Puerto Rico/8/34). The strains were propagated, harvested, and titered following established protocols as previously described^37^. For H5N1, synthetic RNA controls were obtained from Twist Bioscience, consisting of the HA (GenBank: OR051630.1) and NA (GenBank: OR051629.1) segments. All viruses and RNA were stored at −80°C prior to use. These spike-in viruses were selected to avoid background interference, as they are not currently circulating. In contrast, the probe-capture and tiled-amplicon tools were designed using sequences from a broad range of past and current influenza A strains (Section 4.2), supporting their applicability to real-world surveillance.

### 4.2 Design of Genomic Surveillance Tools

The custom IAV-specific probe panel was designed to target 11 different subtypes of IAV: H1N1, H3N1, H5N1, H1N2, H2N2, H3N2, H9N2, H7N9, H4N6, H5N6, and H5N9. These subtypes were selected based on the five most abundant archived IAV sequences in the GISAID database from avian, swine, and human hosts^38^. Additional targets can be incorporated into future versions of the panel as viral evolution and public health priorities change.

Genomes from each subtype were downloaded from GISAID and only filtered for completeness. 525,075 sequences are quality-filtered and clustered by segment at 90% identity, producing a final set of 589 sequences for probe design (See **Table S1** and **Supplementary Methods** for more details on the input genomes). Syotti^39^ was used to design probes with a length of 120 bp (-L 120), a mismatch tolerance of 5 (-d 5), coverage of 90% (-c 0.90), and random input sequence order (-r). This was followed by a fill-gaps command with redesigned probes for gaps larger than 20 bp (-g 20) and a mismatch tolerance of 5 (-d 5). The coverage of the generated probes was validated using Syotti against the spike-in reference genomes (**Supplementary methods**). Probes with alignment lengths greater than 80 bp to the human genome or prokaryotic sequences were removed from the panel. The final probe panel contained 7,448 probes, which were synthesized by Twist Bioscience (San Francisco, California, USA).

Custom tiled amplicon primers were designed for the HA segments of H1N1, H3N2, and H5N1 subtypes using PriMux^22,23^. Briefly, HA sequences were downloaded from GISAID and filtered to remove short sequences (see **Supplementary Methods**). After clustering segments from each subtype at 100% identity with CD-HIT v4.8.1^40^, the reference genomes for the cultured virus strains were added, and multiple sequence alignments were generated for each subtype using MAFFT v7.525^41^. Each alignment was submitted separately to PriMux with options for 450 nt segments with 200 nt overlaps. The resulting primer sets were analyzed with simulate_PCR^42^ with a word-size of 5, maximum of 3 mismatches, and no mismatches allowed within 3 nt of the 3’ end, and visualized in Python v3.9.13 (**Figure S2c**). Subsequent manual curation and iteration were performed to optimize the primer design (see **Supplementary Methods**).

### 4.3 Pre-existing Methods for IAV Sequencing

Primers for whole-segment amplification were adapted from widely used sets for influenza genomic surveillance in clinical samples^7^, referred to herein as “Universal-amplicon”. These primers targeted the conserved non-coding regions at both ends (3’ UTR and 5’ UTR) of each segment. The Comprehensive Viral Research Panel (Twist Biosciences) was adopted to represent an off-the-shelf probe panel, referred to herein as “Probe-Twist”. This panel contains >1 million unique probes targeting 3,153 viral genomes, including influenza A virus genomes from multiple collection dates spanning all four spike-in subtypes (H1N1, H3N1, H3N2, H5N1).

### 4.4 Sample Preparation

#### Direct RNA mixtures of extracted virus RNA and wastewater RNA

Direct RNA mixtures were prepared by separately extracting viral stock RNA and IAV-negative wastewater RNA, then combining them at a series of three targeted concentration ratios to mimic post-extraction conditions (**Figure 1a**). RNA from H1N1, H3N2, and H3N1 viral stocks was isolated using the Direct-zol RNA kit (Zymo Research). Twist synthetic RNA controls for the H5N1 HA and NA segments were used directly without extraction. Wastewater was collected as a grab sample (10/3/2025) from a large city in California, and viral RNA was obtained using the InnovaPrep Concentrating Pipette, followed by extraction with the AllPrep PowerViral Kit (Qiagen) as previously described^18^. Wastewater RNA was treated with DNase (Qiagen). Viral RNA was quantified by dPCR using assays targeting the HA gene and distinguishing the spike-in strains, as previously described^43^. The wastewater RNA was confirmed to be HA-negative for all four IAV spike-in strains, while the dPCR-confirmed concentration of each strain was used to calculate the spike-in volume for preparing virus cocktail at 3 different concentration levels (S1-S3, **Table S10**). Finally, a 5 uL aliquot of each virus cocktail was mixed with 12 uL of wastewater RNA to reach target concentrations of around 5.8, 58.8, and 588 gc/uL per virus in library inputs S1 to S3 (**Figure 1a and Table S6**). These target concentrations were selected based on cultured virus conditions and the detection limits of the dPCR assay; however, concentrations in real wastewater could be lower and more variable^36^. The mass ratio of virus to wastewater background ranged from 4.4×10⁻⁸ to 2.4×10⁻⁶ (S1 to S3, **Table S6**). Virus concentrations in the library inputs were further confirmed by dPCR and closely matched target values for all viruses except for H5N1, which may reflect a one-time experimental error during sample preparation (**Table S10**).

#### Incubation mixtures with spike-in virus in 40-mL wastewater

Incubation samples were prepared by spiking virus stocks into wastewater, followed by 3-hour incubation before concentration and extraction (**Figure 3a**). The incubation time was chosen to approximate the equilibrium of extraviral RNA decay and viral partitioning and adsorption to solids in wastewater35,36. Specifically, 10 µL of each of the three virus stocks (H1N1, H3N1, and H3N2) was spiked into 40 mL aliquots of wastewater influent. Two aliquots were concentrated and extracted immediately, while six aliquots underwent a 3-hour rotation at room temperature before concentration and extraction. Viral RNA was concentrated and extracted from each 40 mL aliquot according to two protocols previously described^43^. Briefly, the Innovaprep method (IP) included the addition of 400µL of 5% Tween 20, centrifugation at 7,000 g, ultrafiltration of the supernatant using the Innovaprep CP Select pipette (CC08004 Unirradiated, InnovaPrep), and elution using elution fluid Tris (InnovaPrep). Nucleic acids were extracted from viral concentrate with the AllPrep PowerViral kit (Qiagen), with an on-column DNase treatment. The Promega method (PMG) utilized the Wizard Enviro Total Nucleic Acid kit (Promega), according to manufacturer’s instructions, followed by RNA purification using Promega RQ1 DNase. For both methods, the final RNA was eluted in 80 µL per sample. The concentrations of all three virus input stocks were tested after same-day extraction using the Direct-zol RNA kit (Zymo Research) and dPCR quantification to avoid extra freeze-thaw cycles.

### 4.5 Library preparation and sequencing

#### Probe capture sequencing using Twist library preparation and enrichment kits

The two probe-capture methods, Probe-IAV and Probe-Twist, followed the Twist Total Nucleic Acids Library Preparation EF Kit 2.0 and the Twist Target Enrichment Standard Hybridization v2 Protocol, with optimizations for viral samples. Briefly, 15 µL of RNA (approximately 50 ng) were used as input for library preparation. First-strand cDNA synthesis was carried out using the ProtoScript II First Strand Synthesis Kit with Random Primer 6 (New England Biolabs) for the Probe-IAV panel, while Superscript IV (Thermo Fisher Scientific) was used for the Probe-Twist panel. For both panels, second-strand cDNA synthesis was performed using the NEBNext Ultra II Non-directional RNA Second Strand Synthesis Module (New England Biolabs). Following lab-specific protocols optimized through independent consultations with technical support, cDNA samples (≤ 25 ng) intended for Probe-IAV were incubated for 5 minutes to achieve a target segment length of 330bp, samples intended for Probe-Twist were incubated for 20 minutes to achieve a segment length of 180-200 bp. After end repair and dA-tailing using NEBNext Ultra II End Prep reagent, adapters were ligated, and each sample was indexed using Twist’s unique dual index system. For barcoding PCR, 12 amplification cycles were used for the Probe-IAV panel, while 10 cycles were performed for Probe-Twist panel.

Following PCR clean-up, 5-8 indexed libraries were multiplexed to a total mass of 1500 ng for each hybridization reaction. Hybridization was performed for 16 hours, followed by washing with magnetic Twist Streptavidin Binding Beads and elution. The post-capture libraries were amplified with 17 cycles in the Probe-IAV and 8 cycles in the Probe-Twist, based on their different panel sizes. Enriched samples were purified before PCR quantification for sample pooling. After the final clean-up, all enriched samples from the customized IAV panel were sequenced on a single NextSeq P2 flow cell (600-cycle 300 PE) together with tiled-amplicon samples, generating approximately 180 Gbp of data. Probe-Twist samples were sequenced separately on a NextSeq P1 flow cell (300-cycle 150 PE). Pooling ratios were decided following the sequencing facilities’ advice, and the resulting sequencing depth for each sample is reported in **Table S7**.

#### Tiled amplicon sequencing

5 µL RNA samples were used as input for HA segment tiled amplicon library preparation. Nuclease-free water served as the negative control, while a mixture of pure virus stock and Twist RNA control was used as the positive control. cDNA was synthesized using the SuperScript IV One-Step Master Mix (Thermo Fisher Scientific). All primers, reactions, and cycling conditions are summarized in **Tables S3-5**. For each sample, four separate reactions were performed using H1 & H3 primer pools 1 and 2, and H5 primer pools 1 and 2, generating 300-415 bp tiled amplicons. The reaction products were purified using AMPure XP beads with a ratio of 1X, and the amplicons were confirmed via e-gel and quantified by Qubit DNA assay. Based on the Qubit concentration, the four PCR products were combined in a 3:3:1:1 mass ratio for H1 & H3 pool 1, H1 & H3 pool 2, H5 pool 1, and H5 pool 2, respectively. Nuclease-free water was added to reach 50 µL per sample as the final input for end-repair and dA-tailing (NEBNext Ultra II module). Following the Twist EF library preparation protocol, Twist adaptors were ligated to the dA-tailed sequences, and each sample was indexed with Twist’s unique dual indices. An additional 8-cycle barcoding PCR was performed. The final enriched libraries, containing ∼400-565 bp segments, were quality-checked using a fragment analyzer, followed by size selection to remove amplicons smaller than 300 bp. Libraries were pooled and sequenced on NextSeq P2 600 cycles 300 PE.

#### Universal amplicon library and Nanopore sequencing

5-µL samples were used for whole-segment universal primer amplicon sequencing library preparation. The same negative and positive controls were used as in the tiled-amplicon method. Primers for whole-segment RT-PCR were taken directly from Zhou *et al.*, 2009^7^. RT-PCR was performed according to the Superscript IV protocol (**Tables S4** and **S5**). After purification with 0.8X beads, amplicons were eluted in 20 µL nuclease-free water. Successful amplification was confirmed by gel electrophoresis, with all samples and the positive control showing detectable bands of varying intensity, while no band was observed in the negative control (**Figure S4**). Up to 200 fmol of purified amplicon was used as input to the Oxford Nanopore ligation library preparation with Native Barcoding Kit 24 V14 (SQK-NBD114.24). Following library preparation, the pooled sample was eluted in 15 µL of elution buffer and quantified by Qubit DNA assay. A total of 50 fmol was loaded onto a minION flowcell (FLO-MIN114) for sequencing. Basecalling was performed using the high accuracy model: dna_r10.4.1_e8.2_400bps_hac@v5.0.0.

### 4.6 Bioinformatic analysis

The reference genomes for four spike-in viruses were downloaded from NCBI using the subtypes and accession numbers provided by the manufacturer or culturing lab. Segments were aligned using MUSCLE (v3.8.31)^44^. The highly conserved 5’ and 3’ UTRs were trimmed from all references prior to mapping to prevent mis-mapping of reads across strains and segments. All raw short reads generated from Illumina sequencing were quality-trimmed using fastp^45^ (v0.24.0) to remove adaptors, polyX tails, and low-quality bases, as well as to filter out reads shorter than 70 bp. For tiled-amplicon sequencing, forward and reverse primers were additionally trimmed using Cutadapt^46^. All high-quality reads were summarized for statistical metrics using Seqkit (v2.4.0)^47^ and mapped to the trimmed reference genomes using Bowtie2 (v2.5.1)^48^. Concordantly mapped paired reads were further filtered with Reformat (BBMap v39.01)^49^ to ensure the total number of indels, deletions, insertions, and substitutions was less than 5. Filtered mapped reads were re-paired and indexed before downstream coverage analysis with Samtools (v1.17)^50^, which summarized mapping statistics, coverage depth, and coverage breadth. Variant calling was performed using Samtools mpileup and iVar (v1.4.2)^51^, with variants filtered by a minimum mapping quality of 20 and a coverage depth of at least 10. BCFtools (v1.17) was also used to generate VCF files and consensus sequences.

### 4.7 Data analysis

The normalization of coverage depth and reads RPKM across different sequencing methods was calculated with equations (1) and (2), respectively.

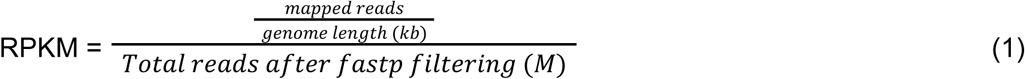

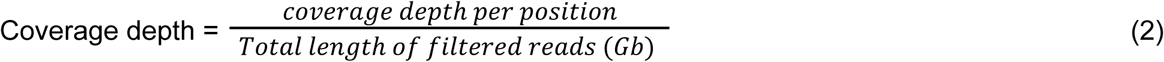

The normality of data was assessed using the Shapiro-Wilk test. If data were normally distributed, statistical differences between different sequencing methods were evaluated using the one-way ANOVA test followed by post hoc pairwise Tukey’s HSD. Otherwise, Kruskal-Wallis tests were applied, followed by the post hoc pairwise Dunn’s test (**Table S12**). All statistical tests were performed using the Python package scipy.stats, and significance was determined at a 95% confidence interval (p < 0.05). All analysis and data visualization were performed with custom python and R scripts.

### 4.8 Economic analysis

The economic analysis includes both cost and labor evaluations for each sequencing method. Costs focus on reagents and consumables, excluding instrument purchase and maintenance. Reagents sold in bulk formats (e.g., 96 reactions) were normalized to a per-sample cost, and sequencing costs were based on the facility and sequencing depth used in this study. Labor time was recorded from manual processing of 24 probe-capture and 57 tiled-amplicon samples. Although labor time could be converted into operational costs, this was not done in the current analysis. All cost data were sourced from purchase records, list prices, or lab documentation and are reported in 2024 U.S. dollars.

### 4.9 Data availability

All raw sequencing data for this project have been deposited in the NCBI sequence Read Archive (SRA) under Bioproject ID: PRJNA1336326. The analysis workflow and reproducible script are available at https://github.com/mj2770/IAV_Seq_WW.

## Author Contributions

MJ: Conceptualization, Methodology, Formal analysis, Investigation, Data Curation, Writing - Original Draft, Writing - Review & Editing, Visualization, Project administration

ALW: Investigation, Writing - Review & Editing JBT: Investigation, Writing - Review & Editing

KLN: Writing - Review & Editing, Supervision, Project administration, Funding acquisition LP: Writing - Review & Editing, Visualization, and Funding acquisition

RSK: Conceptualization, Methodology, Investigation, Data Curation, Writing - Original Draft, Writing - Review & Editing, Visualization, Supervision, Project administration, Funding acquisition

## Acknowledgements

Funding was provided by the UCOP Lab Fees CRT Award (L22CR4507), NIH R00 Award (4R00GM144747-03) and LLNL LDRD 24-SI-006. We thank Dr. Jarno N. Alanko, author of Syotti, for his suggestions during probe design. We are grateful to Anne Beaudoin at EBMUD for assistance with wastewater sample collection. The spike-in virus was cultured by Dr. Staci R. Kane, Dr. Monica K. Borucki, and Jordan Boeck. We prepared sequencing libraries with guidance from Casey Riegler (Twist Bioscience) and Bryan Bach (Functional Genomics Laboratory). Flow-cell loading and sequencing were performed at the Vincent J. Coates Sequencing Laboratory (QB3, UC Berkeley; RRID: SCR_022170) with advice from Shana L. McDevitt and Carrie Rose. Funding to RSK and JBT was provided by the Lawrence Livermore National Laboratory’s Laboratory Directed Research and Development Program. This work was performed in part under the auspices of the U.S. Department of Energy by Lawrence Livermore National Laboratory under Contract DE-AC52-07NA27344, Release Number LLNL-JRNL-2011445. LLNL Disclaimer: This document was prepared as an account of work sponsored by an agency of the United States government. Neither the United States government nor Lawrence Livermore National Security, LLC, nor any of their employees make any warranty, expressed or implied, or assume any legal liability or responsibility for the accuracy, completeness, or usefulness of any information, apparatus, product, or process disclosed, or represent that its use would not infringe privately owned rights. Reference herein to any specific commercial product, process, or service by trade name, trademark, manufacturer, or otherwise does not necessarily constitute or imply its endorsement, recommendation, or favoring by the United States government or Lawrence Livermore National Security, LLC. The views and opinions of authors expressed herein do not necessarily state or reflect those of the United States government or Lawrence Livermore National Security, LLC, and shall not be used for advertising or product endorsement purposes.

## Reference

1. Biggerstaff, M., Cauchemez, S., Reed, C., Gambhir, M. & Finelli, L. Estimates of the reproduction number for seasonal, pandemic, and zoonotic influenza: a systematic review of the literature. BMC Infect. Dis. 14, 480 (2014).

2. Horimoto, T. & Kawaoka, Y. Influenza: lessons from past pandemics, warnings from current incidents. Nat. Rev. Microbiol. 3, 591–600 (2005).

3. Nguyen, T.-Q. et al. Emergence and interstate spread of highly pathogenic avian influenza A(H5N1) in dairy cattle in the United States. Science 388, eadq0900 (2025).

4. CDC. Genetic Sequences of Highly Pathogenic Avian Influenza A(H5N1) Viruses Identified in a Person in Louisiana. Avian Influenza (Bird Flu) https://www.cdc.gov/bird-flu/spotlights/h5n1-response-12232024.html (2025).

5. CDC. Influenza Virus Genome Sequencing and Genetic Characterization. Influenza (Flu) https://www.cdc.gov/flu/php/viruses/genetic-characterization.html (2024).

6. Pagel, C. & Yates, C. A. Tackling the pandemic with (biased) data. Science 374, 403–404 (2021).

7. Zhou, B. et al. Single-reaction genomic amplification accelerates sequencing and vaccine production for classical and Swine origin human influenza a viruses. J. Virol. 83, 10309–10313 (2009).

8. Lee, A. J. et al. Wastewater monitoring of human and avian influenza A viruses in Northern Ireland: a genomic surveillance study. Lancet Microbe 5, 100933 (2024).

9. Khan, M. et al. Significance of wastewater surveillance in detecting the prevalence of SARS-CoV-2 variants and other respiratory viruses in the community – A multi-site evaluation. One Health 16, 100536 (2023).

10. Maqsood, R. et al. Influenza Virus Genomic Surveillance, Arizona, USA, 2023–2024. Viruses 16, 692 (2024).

11. Rothman, J. A. et al. RNA Viromics of Southern California Wastewater and Detection of SARS-CoV-2 Single-Nucleotide Variants. Appl. Environ. Microbiol. 87, e01448–21 (2021).

12. Wyler, E. et al. Pathogen dynamics and discovery of novel viruses and enzymes by deep nucleic acid sequencing of wastewater. Environ. Int. 190, 108875 (2024).

13. Tisza, M. J. et al. Sequencing-Based Detection of Avian Influenza A(H5N1) Virus in Wastewater in Ten Cities. N. Engl. J. Med. 391, 1157–1159 (2024).

14. John, A. et al. Characterizing influenza A virus lineages and clinically relevant mutations through high-coverage wastewater sequencing. Water Res. 287, 124453 (2025).

15. Targeted tiled amplicon based protocol for sequencing the Hemagglutinin (HA) gene segment of seasonal influenza A and influenza B virus from wastewater at high depth of coverage | medRxiv. https://www.medrxiv.org/content/10.1101/2025.10.15.25338105v1.

16. Vo, V. et al. Identification and genome sequencing of an influenza H3N2 variant in wastewater from elementary schools during a surge of influenza A cases in Las Vegas, Nevada. Sci. Total Environ. 872, 162058 (2023).

17. Kantor, R. S., Nelson, K. L., Greenwald, H. D. & Kennedy, L. C. Challenges in Measuring the Recovery of SARS-CoV-2 from Wastewater. Environ. Sci. Technol. 55, 3514–3519 (2021).

18. Jiang, M. et al. Evaluation of the Impact of Concentration and Extraction Methods on the Targeted Sequencing of Human Viruses from Wastewater. Environ. Sci. Technol. 58, 8239–8250 (2024).

19. Kantor, R. S. & Jiang, M. Considerations and Opportunities for Probe Capture Enrichment Sequencing of Emerging Viruses from Wastewater. Environ. Sci. Technol. 58, 8161–8168 (2024).

20. Lail, A. J. et al. Amplicon sequencing of pasteurized retail dairy enables genomic surveillance of H5N1 avian influenza virus in United States cattle. PloS One 20, e0325203 (2025).

21. Kuchinski, K. S., Duan, J., Himsworth, C., Hsiao, W. & Prystajecky, N. A. ProbeTools: designing hybridization probes for targeted genomic sequencing of diverse and hypervariable viral taxa. BMC Genomics 23, 579 (2022).

22. Hysom, D. A. et al. Skip the Alignment: Degenerate, Multiplex Primer and Probe Design Using K-mer Matching Instead of Alignments. PLOS ONE 7, e34560 (2012).

23. Gardner, S. N. et al. Multiplex Degenerate Primer Design for Targeted Whole Genome Amplification of Many Viral Genomes. Adv. Bioinforma. 2014, 101894 (2014).

24. Kent, C. et al. PrimalScheme: open-source community resources for low-cost viral genome sequencing. 2024.12.20.629611 Preprint at 10.1101/2024.12.20.629611 (2024).

25. Wang, M. X., et al. Olivar: towards automated variant aware primer design for multiplex tiled amplicon sequencing of pathogens. Nat. Commun. 15, 6306 (2024).

26. Harrison, K. R., Snead, D., Kilts, A., Ammerman, M. L. & Wigginton, K. R. The Protective Effect of Virus Capsids on RNA and DNA Virus Genomes in Wastewater. Environ. Sci. Technol. 57, 13757–13766 (2023).

27. Nash, D. et al. A Genome Sequence Variant Monitoring Program for Seasonal Influenza A H3N2 and Respiratory Syncytial Virus A using Wastewater-Based Surveillance in Ontario, Canada. 2025.07.29.667219 Preprint at 10.1101/2025.07.29.667219 (2025).

28. Davis, J. J. et al. Analysis of the ARTIC Version 3 and Version 4 SARS-CoV-2 Primers and Their Impact on the Detection of the G142D Amino Acid Substitution in the Spike Protein. Microbiol. Spectr. 9, e01803–21.

29. Chakraborty, C., Das, A., Bhattacharya, M. & Islam, Md. A. Mutations in the influenza virus, primarily H5N1, enhance the virus’s virulence, favor receptor interaction, and increase drug resistance. Int. J. Surg. Lond. Engl. 111, 4128–4131 (2025).

30. Ping, J. et al. PB2 and Hemagglutinin Mutations Are Major Determinants of Host Range and Virulence in Mouse-Adapted Influenza A Virus. J. Virol. 84, 10606–10618 (2010).

31. McCall, C. et al. Targeted Metagenomic Sequencing for Detection of Vertebrate Viruses in Wastewater for Public Health Surveillance. ACS EST Water 3, 2955–2965 (2023).

32. The Effects of mismatches on DNA Capture by Hybridization | Twist Bioscience. https://www.twistbioscience.com/resources/white-paper/effects-mismatches-dna-capture-hybridization.

33. Kim, H., Webster, R. G. & Webby, R. J. Influenza Virus: Dealing with a Drifting and Shifting Pathogen. Viral Immunol. 31, 174–183 (2018).

34. Karthikeyan, S. et al. Wastewater sequencing reveals early cryptic SARS-CoV-2 variant transmission. Nature 609, 101–108 (2022).

35. Wang, A. L. et al. Benchmarking concentration and direct extraction methods for wastewater-based surveillance of eight human respiratory viruses: implications for rapid application to novel pathogens. 2024.11.27.625007 Preprint at 10.1101/2024.11.27.625007 (2024).

36. Wolfe, M. K. et al. Wastewater-Based Detection of Two Influenza Outbreaks. Environ. Sci. Technol. Lett. 9, 687–692 (2022).

37. Ye, Y., Ellenberg, R. M., Graham, K. E. & Wigginton, K. R. Survivability, Partitioning, and Recovery of Enveloped Viruses in Untreated Municipal Wastewater. Environ. Sci. Technol. 50, 5077–5085 (2016).

38. Daniels, R. S. & McCauley, J. W. The health of influenza surveillance and pandemic preparedness in the wake of the COVID-19 pandemic. J. Gen. Virol. 104, 001822 (2023).

39. Alanko, J. N. et al. Syotti: scalable bait design for DNA enrichment. Bioinformatics 38, i177–i184 (2022).

40. Fu, L., Niu, B., Zhu, Z., Wu, S. & Li, W. CD-HIT: accelerated for clustering the next-generation sequencing data. Bioinformatics 28, 3150–3152 (2012).

41. Katoh, K., Misawa, K., Kuma, K. & Miyata, T. MAFFT: a novel method for rapid multiple sequence alignment based on fast Fourier transform. Nucleic Acids Res. 30, 3059–3066 (2002).

42. Gardner, S. N. & Slezak, T. Simulate_PCR for amplicon prediction and annotation from multiplex, degenerate primers and probes. BMC Bioinformatics 15, 237 (2014).

43. Wang, A. L. et al. Benchmarking Concentration and Extraction Methods for Wastewater-Based Surveillance of Eight Human Respiratory Viruses: Implications for Rapid Application to Novel Pathogens. Environ. Sci. Technol. 10.1021/acs.est.4c13635 (2025) doi:10.1021/acs.est.4c13635.

44. Edgar, R. C. MUSCLE: a multiple sequence alignment method with reduced time and space complexity. BMC Bioinformatics 5, 113 (2004).

45. Chen, S., Zhou, Y., Chen, Y. & Gu, J. fastp: an ultra-fast all-in-one FASTQ preprocessor. Bioinformatics 34, i884–i890 (2018).

46. Martin, M. Cutadapt removes adapter sequences from high-throughput sequencing reads. EMBnet.journal 17, 10–12 (2011).

47. Shen, W., Le, S., Li, Y. & Hu, F. SeqKit: A Cross-Platform and Ultrafast Toolkit for FASTA/Q File Manipulation. PLOS ONE 11, e0163962 (2016).

48. Langmead, B. & Salzberg, S. L. Fast gapped-read alignment with Bowtie 2. Nat. Methods 9, 357–359 (2012).

49. Bushnell, B. BBMap: A Fast, Accurate, Splice-Aware Aligner. https://www.osti.gov/biblio/1241166 (2014).

50. Li, H. et al. The Sequence Alignment/Map format and SAMtools. Bioinformatics 25, 2078–2079 (2009).

51. Grubaugh, N. D. et al. An amplicon-based sequencing framework for accurately measuring intrahost virus diversity using PrimalSeq and iVar. Genome Biol. 20, 8 (2019).

