## Supplemental Materials for "Developing and Benchmarking One Health Genomic Surveillance Tools for Influenza A Virus in Wastewater"

**Tool for Influenza A Virus in Wastewater**

Minxi Jiang^a^, Audrey L.W. Wang^a^, James B. Thissen^b^, Kara L. Nelson^a^, Lenore Pipes^c^*, Rose S. Kantor^b^*

a. Department of Civil and Environmental Engineering, University of California, Berkeley, CA, USA
b. Physical and Life Sciences Directorate, Lawrence Livermore National Laboratory, Livermore, CA, USA.
c. Pacific Biosciences Research Center, University of Hawai’i at Mānoa, Honolulu, HI, USA

**Contents:**

**Supplementary Tables: 14**

**Supplementary Figures: 4**

**Supplementary Methods**

**Supplementary Tables**

**Table S1.** Metadata for sequences included in the probe design, including collection date, download date, virus strains, virus hosts, and the number of sequences per strain per segment before and after data clean-up

**Table S2.** Probe design parameters and resulting probe numbers during the probe optimization

**Table S3.** Tiled-amplicon primer sequences designed in this study

**Table S4.** Amplicon-based method PCR reaction recipe

**Table S5.** Amplicon-based method PCR thermal cycling conditions

**Table S6.** Targeted and dPCR-quantified HA segment concentrations for each virus across direct mixtures S1–S3

**Table S7.** Averaged sequencing depth across triplicates for each sample

**Table S8**. Averaged log10 (RPKM) and coverage breadth of all segments across all sequencing methods and all virus strains

**Table S9.** The slope and R^2^ value of the quantitative correlation between log10 RPKM and log10 dPCR concentration of HA

**Table S10**. dPCR concentration and partitioning of the extracted virus stock, wastewater, Twist H5N1 standards, and S1-S3 sample mixtures

**Table S11**. dPCR-measured viral RNA concentrations in incubation samples after extraction of two concentration and extraction methods (IP v.s. PMG)

**Table S12.** Statistical analysis on coverage breadth, dPCR conc., and RPKM from each strain under each sequencing method, comparing concentration and extraction methods of Innovaprep + Powerviral (IP) with Promega (PMG)

**Table S13.** Log2 fold change of coverage breadth of all IAV strains’ 8 segments from two wastewater processing methods and four sequencing methods

**Table S14.** Economic evaluation of sequencing methods based on capital costs and labor hours

**Supplementary Figures**

**Figure S1.** Host distribution of IAV genomes included in the probe design

**Figure S2.** Predicted coverage depth and breadth from the Probe-IAV and Tiled-amplicon panel

**Figure S3.** Comparison of the coverage breadth and depth for all IAV segments from Probe-IAV, Probe-Twist, and Universal-amplicon

**Figure S4**. Gel electrophoresis results of amplicons enriched using the universal primers prior to Nanopore sequencing library preparation

### **Supplementary Methods**

### Design and validation of customized probes and primers

- Library preparation for each sequencing method

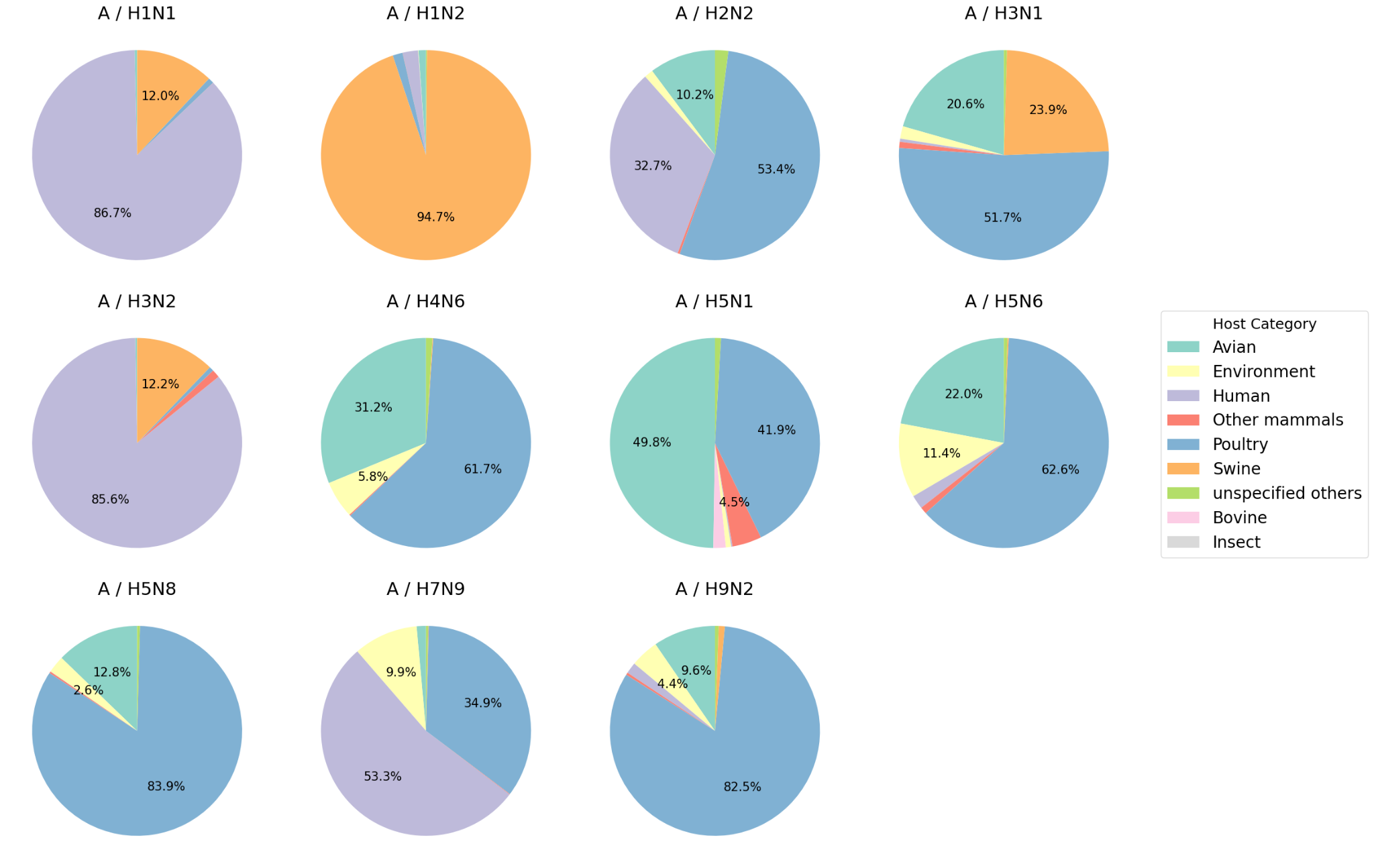

**Figure S1.** Host distribution of IAV genomes included in the probes and primers design

Further filters were applied to select high-quality genomes by segment using a customized Python script. Sequences with ambiguous bases and those with lengths more than 100 bp above or below the median segment length were discarded. All high-quality sequences were clustered by segment with 90% identity using CD-Hit, resulting in a total of 563 sequences among 8 segments and 11 subtypes (**Table S1)**. Twenty-six additional sequences from four spike-in viruses (8 segments each for H1N1, H3N1, H3N2, and only HA and NA segments for H5N1) were added, resulting in a final input of 589 sequences for probe design (**Table S1**).

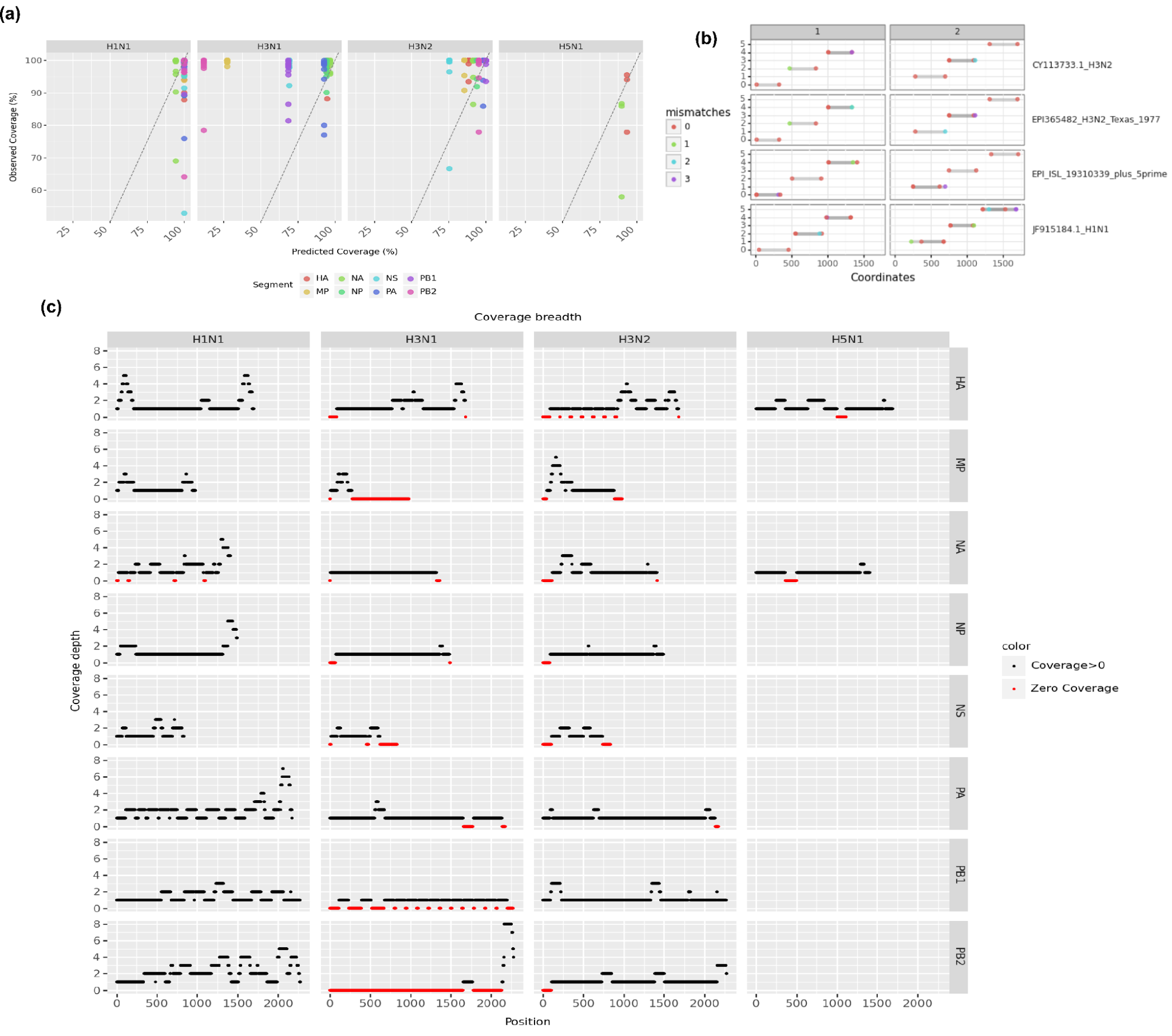

**Figure S2.** Predicted performance regarding coverage depth and breadth from the two customized sequencing methods (Probe-IAV and Tiled-amplicon). **(a**) Predicted v.s. Actual coverage breadth (%) of all segments from all strains by Probe-IAV. **(b)** Simulated PCR results of designed primers; **(c)** Predicted coverage depth of all segments across genomes by Probe-IAV.

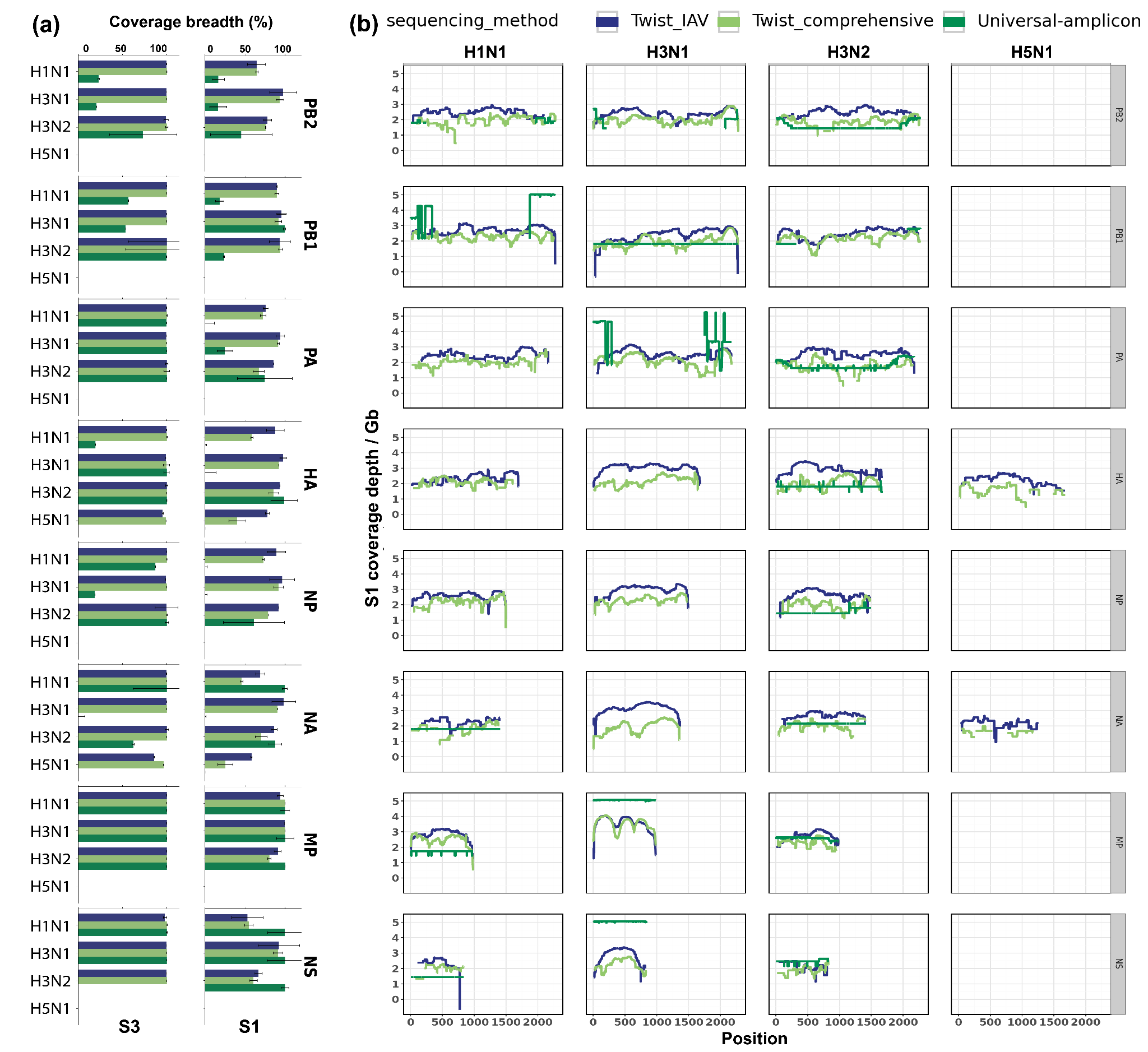

**Figure S3**: Comparison of the coverage breadth and depth for all IAV segments from the three whole-genome sequencing methods (Probe-IAV, Probe-Twist, and Universal-amplicon). **(a)** Coverage breadth of all segments of H1N1, H3N1, H3N2, and H5N1 in sample S1 and S3; **(b)** Coverage depth across the genome of all 8 segments of H1N1, H3N1, H3N2, and H5N1 in sample S1.

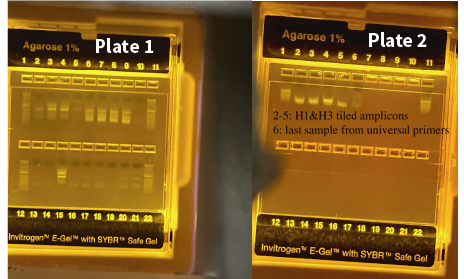

| Plate | Well | Samples | Bands |
| --- | --- | --- | --- |
| Plate 1 | 1, 11, 12, and 22 | Gradient standards | Yes |
| Plate 1 | 10, 13, and 14 | Sample S1 triplicates | Yes |
| Plate 1 | 16, 17, and 18 | Sample S2 triplicates | Yes |
| Plate 1 & Plate 2 | 20, 21 on Plate 1 and 6 on Plate 2 | Sample S3 triplicates | Yes |
| Plate 1 | 2,3, 4, and 5 | IP samples at time 0 and 3 hours with spike-in viruses | Yes |
| Plate 1 | 6,7,8,and 9 | PMG spike-in samples at time 0 and 3 hours | Yes |
| Plate 1 | 19 | Negative control | No |
| Plate 1 | 15 | Positive control containing extracted H3N2 RNA (~1.6 × 10⁵ gc/µL) as quantified by dPCR | Yes |

**Figure S4.** Gel electrophoresis results of amplicons enriched using the universal primers prior to Nanopore sequencing library preparation. The DNA ladder is designed for sizing dsDNA in the range of 100 bp to 15,000 bp, with a reference band at 1,500 bp for orientation. Universal primer–enriched amplicons are expected to be 900-2,500 bp. Detailed well assignments, corresponding samples, and band information are provided in the table below the figure.

### **Supplementary Methods**

### 1. **Design and validation of customized probes and primers**

### Probe-IAV panel

The IAV-specific probe panel was designed to target 11 different subtypes of IAV genomes: H1N1, H3N1, H5N1, H1N2, H2N2, H3N2, H9N2, H7N9, H4N6, H5N6, and H5N9. These subtypes were selected based on the five most abundant archived IAV sequences in the GISAID database from avian, swine, and human hosts ^1^. Genome sequences for each subtype were downloaded from GISAID on 6/7/2024, including only complete genomes with all 8 segments. To balance the representation of different subtypes, specific time frames were applied for downloading sequences (**Table S1**). Metadata of those downloaded sequences revealed that the one-year data for H1N1 and H3N2 predominantly comprised sequences from human hosts (**Figure S1**). To ensure more diverse host representation, additional H1N1 and H3N2 sequences from avian and swine hosts, collected within the past 10 years (5/15/2014 to 5/15/2024), were manually included. This process resulted in a final set of 525,075 IAV genomes, and diverse hosts from avian, swine, and human (**Table S1**).

Syotti was selected for probe design due to its efficiency in generating minimal probes in a short time ^2^. Different design parameters were tested to balance probe efficiency and total amount, which includes mismatch number, genome coverage, and coverage gaps (**Table S2**). The final set was designed with a probe length of 120 bp (-L 120), a mismatch tolerance of 5 (-d 5), coverage of 90% (-c 0.90), and random order (-r). This was followed by a fill-gaps command with redesigned probes for gaps larger than 20 bp (-g 20) and a mismatch tolerance of 5 (-d 5). The coverage of the designed probes was validated using Syotti against the spike-in reference genomes (**Figure S2a and S2b**). Notably, several segments of the H3N1 genome—particularly PB1, PB2, and MP—showed low predicted coverage (**Figure S2c**; NS: 69.21%, PB1: 68.6%, MP: 27.49%, PB2: 11.71%). However, experimental results showed that actual coverage exceeded 95% in nearly all cases, with only one PB2 and two PB1 samples falling slightly lower (between 80–90%). We also observed that certain regions—particularly the ends of genome segments—which may be attributed to discrepancies in segment lengths among sequences deposited in public databases. The final probe panel was also checked against human genome background data (GRCh38.p14: RefSeq GCF_000001405.40 and CHM13v2.0: RefSeq GCF_009914755.1) and the nt_prok NCBI database (accessed on Jun 5th, 2024). Probes with alignment lengths greater than 80 bp to these references were filtered out from the panel.

### Tiled-amplicon panel

The customized tiled-amplicon panel was designed for the HA segment from H1N1, H3N2, and H5N1. Primers were designed using PriMux (Hysom 2012, Gardner 2014). Reference sequences downloaded from GISAID included HA segments from H3N2 and H1N1 genomes collected between May 15, 2023 - May 15, 2024 and H5N1 genomes collected between May 15, 2019 - May 15, 2024. Sequences were filtered to remove sequences with >5% Ns, trimmed to remove 5’ and 3’ terminal Ns, and filtered to remove sequences longer or shorter than the median sequence length by more than 100 nt. After clustering the filtered sequences at 100% identity within each subtype, the reference genomes for the cultured virus strains were added to the resulting files for H1N1, H3N2, and H5N1, respectively. Multiple sequence alignments were generated with MAFFT v7.525 for each of the three HA subtypes and submitted separately to PriMux with settings indicating 450 nt amplicons with 200 nt overlaps. The resulting primer sets were split into pools 1 and 2, to produce non-overlapping amplicons, and primer pools were analyzed within each subtype using simulate_PCR ^3^ with a word-size of 5, maximum of 3 mismatches, and no mismatches allowed within 3 nt of the 3’ end. Outputs from simulate_pcr were visualized in Python v3.9.13 (**Figure S2c**). Where multiple primer sets were produced for a given tile, the best set was chosen based on the length of the amplicon produced and mismatches to the cultured strain(s) of that subtype. Primers were searched within the multiple sequence alignments using Geneious Prime v2024.0.5 to visualize the representativeness of each primer across all references. Additional degenerate nucleotides were added where needed (**Table S3**). Potential primer dimers and self-dimers were assessed with the ThermoFisher Multiple Primer Analyzer) and iteratively removed by choosing alternative primers from the sets produced by PriMux. Final primers were divided into two pools and reanalyzed with simulate_pcr and the Multiple Primer Analyzer. Primer dimers necessitated manual redesign of four primers (**Table S3**). This was performed by extracting the appropriate regions of the consensus sequence from the multiple sequence alignment for input into Primer3Plus (https://www.primer3plus.com/index.html; Untergasser 2012). Finally, a single short amplicon (228 bp) was predicted to be produced from H1N1 by primers targeting H1 and H5. Thus we conducted RT-PCR for H5 separately from the combined H1 and H3 reactions for a total of 4 reactions (2 pools for each).

To validate the primer design, each tile was individually amplified using RNA extracted from the benchmarking strains. PCR products were run on 1% agarose E-Gels with SYBR Safe DNA stain to confirm amplicon presence. Samples without clear bands were either re-amplified or redesigned. In the initial trial (0.5 nM primer concentration, 25-µL reactions, 55 °C annealing, as recommended for the Invitrogen SuperScript IV One-Step RT-PCR System), 9 tiles failed. Four were redesigned with higher degeneracy, requiring higher primer concentrations and lower annealing temperatures. After optimization (2.5 nM primers, 50-µL reactions, 50 °C annealing), 7 of the previously failed tiles amplified successfully, while H1 tile 5 and H5 tile 0 still failed. A final redesign step was then performed, removing degeneracy to increase primer specificity.

2. **Library preparation for each sequencing method**

### Probe capture sequencing using Twist library preparation and enrichment kits

The two probe-capture methods include Probe-IAV and Probe-Twist, both follow the Twist Total Nucleic Acids Library Preparation EF Kit 2.0 and the Twist Target Enrichment Standard Hybridization v2 Protocol, with optimizations for virus samples. Briefly, 15 µL of the prepared samples (with approximately 50 ng RNA) were used as input for library preparation. First-strand cDNA synthesis was carried out using the ProtoScript II First Strand Synthesis Kit with Random Primer 6 for the customized IAV panel, while Superscript IV was used for the Probe-Twist panel. Second-strand cDNA synthesis was performed using the NEBNext Ultra II Non-directional RNA Second Strand Synthesis Module for both panels. For fragmentation, each cDNA sample (≤ 25 ng) was incubated for 5 minutes in the Probe-IAV panel to achieve a target segment length of 330 bp, while a 20-minute incubation in the Probe-Twist panel resulted in a segment length of 180-200 bp. After end repair and dA-tailing, adapters were ligated, and each sample was indexed using Twist’s unique dual index system. For barcoding PCR, 12 amplification cycles were used in the Probe-IAV panel, while only 10 cycles were performed in Twist’s probe panel.

Following PCR clean-up, 5-8 indexed libraries were multiplexed to a total mass of 1500 ng for each hybridization reaction. Hybridization was performed for 16 hours, followed by washing with magnetic Twist Streptavidin Binding Beads and elution. The post-capture libraries were amplified with 17 cycles in the Probes_IAV and 8 cycles in the Probe-Twist, based on their different panel sizes. Enriched samples were purified before PCR quantification for sample pooling. After the final clean-up, all enriched samples from the customized IAV panel were sequenced on a NextSeq P2 600-cycle 300 PE, and Probe-Twist samples were sequenced on NextSeq P1 300-cycle 150 PE.

### Tiled-amplicon sequencing

5 µL samples were used as input for H-gene tiled-amplicon sequencing library preparation. Nuclease-free water served as the negative control, while a mixture of pure virus stock and Twist RNA control was used as the positive control. cDNA was synthesized using the SuperScript IV One-Step Master Mix (Thermo Fisher Scientific). All primers, reactions, and cycling conditions are summarized in **Table S4 and Table S5**. For each sample, four separate reactions were performed using H1 & H3 primer pools 1 and 2, and H5 primer pools 1 and 2, generating 300-415 bp tiled-amplicons. The reaction products were purified using 1X beads, and the amplicons were confirmed via e-gel and quantified by Qubit DNA assay. Based on the Qubit concentration, the four PCR products were combined in a mass ratio of 300 ng:300 ng:100 ng:100 ng, and nuclease-free water was added to reach 50 µL per sample as the final input for end-repair and dA-tailing (NEBNext Ultra II module). Following the Twist EF library preparation protocol, Twist adaptors were ligated to the dA-tailed sequences, and each sample was indexed with Twist’s unique dual indices. An additional 8-cycle barcoding PCR was performed to minimize PCR errors. The final enriched libraries, containing ~400-565 bp segments, were quality-checked using a fragment analyzer, followed by an optional size selection to remove amplicons smaller than 300 bp. All qualified libraries were pooled and sequenced on Nextseq P2 600 cycles 300 PE.

### Universal amplicon library and Nanopore sequencing

5 µL samples were used for whole-segment universal primer amplicon sequencing library preparation. The same negative and positive controls were used as in tiled-amplicon sequencing. Primers for the whole-segment RT-PCR were drawn directly from Zhou 2009. RT-PCR was conducted according to the protocol for Superscript IV (**Table S4 and Table S5**). After purification with 0.8X beads, amplicons were eluted in 20 µL. Up to 200 fmol of purified amplicon was used as input to the Oxford Nanopore ligation library preparation with Native Barcoding Kit 24 V14 (SQK-NBD114.24). Following library preparation, the pooled sample was eluted in 15 µL of elution buffer and quantified by Qubit DNA assay. A total of 50 fmol was loaded onto a minION flowcell (FLO-MIN114) for sequencing. Basecalling was performed using the high accuracy model: dna_r10.4.1_e8.2_400bps_hac@v5.0.0.

1.	Daniels, R. S. & McCauley, J. W. The health of influenza surveillance and pandemic preparedness in the wake of the COVID-19 pandemic. *J. Gen. Virol.* **104**, 001822 (2023).

2.	Alanko, J. N. *et al.* Syotti: scalable bait design for DNA enrichment. *Bioinformatics* **38**, i177–i184 (2022).

3.	Gardner, S. N. & Slezak, T. Simulate_PCR for amplicon prediction and annotation from multiplex, degenerate primers and probes. *BMC Bioinformatics* **15**, 237 (2014).
